# Simulation-based Bayesian deep learning enables uncertainty-aware tumor fraction estimation in cell-free DNA

**DOI:** 10.64898/2026.06.15.732265

**Authors:** Hadas Volkov, Maria Raitses-Gurevich, Meitar Grad, Rani Shlayem, Artem Danilevsky, Tami Rubinek, Malka Gorfine, Noam Shomron

## Abstract

**Background:** Estimating tumor fraction from whole-genome cell-free DNA sequencing is critical for liquid biopsy, but is hampered by weak signals and baseline noise at low tumor fractions. Existing computational methods often require matched controls or large labeled datasets for training and lack uncertainty quantification. To address these gaps, we developed purNPE, a Bayesian deep-learning framework trained without labeled cancer cell-free DNA samples. Specifically, purNPE leverages a two-part generative model: one component simulates diverse tumor copy-number profiles based on evolutionary genealogies, while a second, data-driven component learns and replicates realistic sequencing background patterns from cancer-free cell-free DNA. By training a Neural Posterior Estimator on synthetic tumor profiles augmented with learned noise, purNPE performs amortized inference in milliseconds without needing a reference sample set at inference.

**Results:** In a real-world pan-cancer cohort, purNPE achieved comparable performance with existing methods against orthogonal mutant-allele-fraction validation (MAE **= 0.066**). In silico and semi-synthetic experiments suggested analytical sensitivity around 1% tumor fraction under the evaluated conditions and showed strong classification accuracy in low tumor fractions (AUC **= 0.98** for TF ***≤* 3%** versus controls).

**Conclusions:** This work provides a framework for using simulation-based inference to derive calibrated, uncertainty-aware TF estimates, offering a potential alternative to traditional data-dependent methods.

## 1 Background

The identification of circulating tumor DNA (ctDNA) from plasma is a cornerstone of liquid biopsy, a non-invasive approach that offers a window into tumor genomics, facilitating the monitoring of treatment response, recurrence risk stratification, and early cancer diagnosis [1–4]. A fundamental task is to accurately quantify the tumor fraction (TF) of ctDNA from mixed cell-free DNA (cfDNA) sequencing reads. Many clinically relevant scenarios operate in the low single-digit TF range, where signal is present but weak (*<* 5%) [5–7]. In this regime, ctDNA signals are substantially diluted and confounded by technical noise, making accurate TF estimation challenging [8].

To meet this challenge, the field has developed multiple analytical strategies that interrogate short-read sequencing data acquired by Next Generation Sequencing (NGS). Among commonly used strategies are methods that aim to infer TF from the variant allele frequencies (VAFs) of known somatic mutations [9]. Nonetheless, these approaches have several limitations obscuring their reliability and usefulness in changing scenarios. There is a need for prior knowledge regarding the credibility of the somatic mutation, acquired either by tumor-informed variants or carefully curated hotspot panels [1]. Secondly, VAF can be misinterpreted in cases where the genomic region is either affected by a copy number variation (CNV) that alters allele composition or in events where genetic variability changes due to the presence of different subclones [10]. Fragmentomics, the analysis of statistical variation observed in fragment features from ctDNA, provides complementary signals that can be used for TF estimation. However, their utility for precise TF estimation varies by context and cancer type, and depends on coverage and assay design [3, 11–14].

The general-purpose methodology of CNV analysis, utilizing one of the hallmarks of cancer for the assessment of cancer purity levels in sequencing data, has long been proven efficient. Cancer is known for high levels of focal CNVs and aneuploidy variations and their contribution to tumorigenesis, with roughly 30% of the cancer genome affected by such events [15]. In solid tumors, around 25% of the genome is altered by big whole-arm or whole-chromosome gains and losses [16]. This statistical variation phenomenon is being exploited in the development of maximum likelihood estimation methods for the detection of purity levels from tumor sequencing and subsequently being adapted by similar approaches for the assessment of TF in cfDNA sequencing [7, 17–23]. Multiple methods with tractable likelihood have been proposed over the years, each differing in the underlying assumptions, the specific type of sequencing assay for which they are relevant, and the resulting estimations. Nonetheless, a commonality of these methods is their simplification of the aberration landscape in a way that allows reasonable computing efforts. Secondly, traditional methods operate by segmentation algorithms or hidden Markov models processing a single genome at a time, meaning they are incapable of learning from past examples. Lastly, the need for matched dedicated control samples, or a pool of reference samples, for the reduction of experiment-specific sequencing noise from the target sample is a significant hurdle, drastically tempering the results if not provided [24].

With the rise of machine learning (ML), and in particular deep learning (DL), many promising methods have emerged for accurate TF estimation from data [11, 25]. These approaches often sidestep the need to specify an explicit likelihood and thereby provide the flexibility to model complex, nonlinear relationships. Nevertheless, a major limitation governing the supervised training regime, the most common and effective approach, is the need for vast, accurately labeled, and diverse datasets for training. When sufficient sample sizes are not provided, generalizability is compromised, and transferability among assays, sequencing depth, and cancer types becomes questionable. Obtaining large, diverse, well-labeled datasets is challenging; domain shift across assays, depths, and cancer types often limits generalization.

A potential path to overcome data scarcity is realism-driven simulation [26–30]. A common simplifying view is that cancer evolution can be approximated by a natural coalescent process with roughly constant growth rate and episodic selective sweeps that reshape the tumor’s Copy Number Profile (CNP) and clonal composition [31, 32]. Such genealogical tree processes can be adapted into generative engines that produce CNPs at scale. Here, we propose using CNP simulations within a Simulation-Based Inference (SBI) framework. We argue that simulated and real CNPs are vast and structured, resulting in a likelihood term that is intractable. Neural Posterior Estimation (NPE), an advanced SBI approach, addresses this by bypassing explicit likelihood computation and replacing it with an approximation of the posterior distribution with neural networks [33, 34]. This strategy overcomes the need for large, labeled datasets; instead, it leverages simulations to generate virtually unlimited synthetic data with known ground-truth.

In this study, we developed purNPE, a likelihood-free Bayesian framework that applies NPE to quantify TF and tumor ploidy levels from cfDNA sequencing data. Our approach is composed of two decoupled elements: a noise generator modeled from existing sequencing data to reproduce global variability patterns observed in cfDNA sequencing along the genome, and a CNP simulator that reproduces the somatic evolution of tumor genomes. The posterior estimator is trained without labeled cancer cfDNA samples: tumor copy-number signals are simulated, whereas sequencing-noise structure is learned from cancer-free cfDNA controls through the Generative Noise Model (GNM). The computational cost of simulation and training is paid once upfront; thereafter, inference on new samples is amortized, requiring only a fast forward pass through the trained network. As such, purNPE delivers calibrated, probabilistic TF estimates in milliseconds and does not require a matched control sample or cohort-specific reference panel at inference.

The study makes four methodological contributions. First, it adapts simulation-based neural posterior estimation to joint TF and ploidy inference from cfDNA CNPs. Second, it introduces a data-driven GNM for augmenting clean simulated tumor signals with realistic cfDNA sequencing artifacts. Third, it returns calibrated posterior distributions rather than point estimates alone, enabling uncertainty-aware interpretation in low-signal regimes. Fourth, it evaluates the approach using simulation diagnostics, semi-synthetic spike-ins, a public pan-cancer cfDNA cohort with an orthogonal Mutant Allele Fraction (MAF) proxy, and independent clinical cohorts, with software and model resources provided for reuse.

## 2 Methods

### 2.1 Tumor evolutionary histories

We aim at generating a broad library of biologically plausible tumor evolutionary histories covering major Copy Number Aberration (CNA) archetypes that will act as the clean CNPs basis. For this, we designed a simulation strategy leveraging the CNAsim single-cell simulator [27]. Our approach was centered around creating a diverse set of plausible cancer genome landscapes, aiming at capturing a wide spectrum of genomic instability observed in clinical tumors. This process involved three main steps, defining biological archetypes, randomizing parameters within each archetype, and assembling the final simulation command.

We first defined five distinct evolutionary archetypes to serve as templates. These archetypes were designed to represent patterns of tumor evolution and genomic instability and plausible across major CNA modes. (1) Diploid quiet represents tumors with mostly diploid genome and minimal CNAs. (2) Aneuploid without Whole Genome Doubling (WGD) (chromosome instability (CIN) dominant), models diploid tumors characterized by a high rate of whole-chromosome or chromosome-arm level events, widespread aneuploidy. (3) Diploid with focal instability, representing tumors with a high rate of focal CNAs but potentially lower levels of whole-CIN. (4) WGD stable, tumors that undergo an early WGD event but subsequently evolve with relative genomic stability. (5) WGD unstable, most chaotic genomes, WGD event is followed by ongoing high rates of focal and CIN.

We proceeded with 1,000 evolutionary history simulations; an archetype was selected based on a weighted probability. Then, key CNAsim parameters were randomized within biologically plausible ranges tailored to that specific archetype. More specifically, we shape clonal structure by tuning selective sweeps and event placement (e.g., from 0-1 for “diploid quiet” to 3-6 for “WGD unstable”). CNA event rates are the rates of focal CNAs (e.g., founder-event-mult, placement-param) and chromosome-level events (e.g., chrom-rate-founder, chrom-rate-clone) were adjusted. For example, “diploid focal” histories were assigned high focal rates, while “WGD unstable” histories were given high rates for focal and chromosomal events. CNA characteristics are universal parameters as the mean size of focal CNAs and the ratio of amplifications to deletions was randomized for every simulation for further variability.

The selected archetype and its randomized parameters were used to construct a unique command-line instruction for CNAsim. Each command specified the full evolutionary model for one tumor, including the number of cells to simulate (n = 500), the potential for a WGD event, the complete set of event rates, and the approximate clonal architecture.

### 2.2 Bulk TF adjusted CNPs formation

Following the generation of the evolutionary histories, we aim at creating a large-scale dataset of 250,000 clean, bulk CNPs with a ground truth TF. Each bulk profile was generated through a multi-step stochastic process; we first determine the TF by randomly sampling a Beta distribution (*α* = 1.001, *β* = 7.0). This parameterization creates a distribution heavily skewed towards low TF values (e.g., *<* 15%), ensuring that the model would be mostly trained on the clinically challenging low-TF regime, while still allowing high TF simulations to occur in low probability. This, in turn, defined the total number of tumor cells and normal cells in the final simulated mixture. Subclonal composition to enlarge intra-tumor heterogeneity was determined by a random draw from a Dirichlet distribution by the relative proportions of each sub-clone present in the specific history. The concentration parameter (*α*) was drawn from a Gamma distribution (shape = 0.5, scale = 0.3). This hierarchical approach ensures a wide variety of subclonal architectures, from tumors dominated by a single clone to a balanced, polyclonal mixture. Based on these sampled proportions, a “bag of cells” was created by performing multinomial sampling to determine the exact count of cells drawn from each subclone, followed by randomly selecting the corresponding individual cells from the CNAsim output.

Finally, bulk CNP was constructed by taking a weighted average of the bulk tumor profile and a diploid profile (i.e., absolute copy number of 2 at all loci). The weights were determined by the tumor TF sampled in the first step.

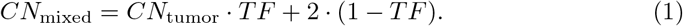

The mixed absolute CNP was converted into a log_2_-transformed copy ratio relative to a diploid baseline. This final vector of log_2_ values represented the “clean signal” for a single simulation. For joint posterior training, the ploidy label was defined as the genome-wide, bin-weighted mean absolute copy number of the sampled tumor component before dilution with diploid cfDNA; TF was the mixture proportion applied during mixing. It is important to note that no normalization of the log_2_ signal was performed, as is common in other bioinformatic pipelines. This was done with the direct intention of allowing the global amplitude to be learned by the subsequent neural network.

### 2.3 Construction of GNM and noise sampling

A data-driven GNM was designed to learn and subsequently reproduce on-the-fly the complex, structured trends and artifacts inherent in real cfDNA sequencing data. The model was trained using different cancer-free control batches according to the evaluation setting, with held-out controls excluded whenever they served as target backgrounds or leakage-sensitive validation data. In general, the available batches were cancer-free control cfDNA samples from three previously published pan-cancer cfDNA WGS datasets (EGAD00001005339 (n = 30), EGAD00001006237 (n = 30) and EGAD00001006132 (n = 24)), and the additional breast cancer (n = 9) and Pancreatic Ductal Adenocarcinoma (PDAC) control samples, where the PDAC samples were divided into two batches according to the hospital of origin (n = 6 each). The specific exclusion policy for each benchmark is stated in the corresponding Results section.

For each sample, the GC- and mappability-corrected log_2_ read-depth profile across a predefined whitelist set of 500-kb autosomal genomic bins was processed (GRCh38 coordinates; 4,784 bins after filtering). The profiles were robustly Z-scored by subtracting the per-sample median and dividing by the median absolute deviation (MAD) to normalize for sample-specific variance:

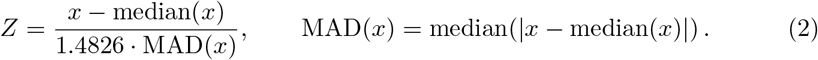

Any remaining missing values post-normalization were imputed with the per-bin median per sample. To ensure the model learned from a representative batch of control profiles, we identified and removed outlier samples within the batch. This was achieved by modeling the relationship between the profile standard deviation and sequencing depth and removing samples with large residuals from this fit (greater than 5 times the MAD of the residuals).

Per-batch modelling was then performed using a Principal Component Analysis - Gaussian Mixture Model (PCA-GMM) [35] approach. First, PCA was applied to the Z-scored profiles of a given batch to identify the dominant components of variation, with the number of principal components (PCs) chosen to explain 95% of the variance. The high-dimensional Z-scored data is projected into a lower-dimensional latent space (*S*) using the PCs.

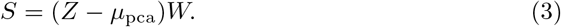

where *µ*_pca_ is the mean vector of the training data and W is the projection matrix containing the top PCs. Next, a GMM with a diagonal covariance is fitted to the data in PC space, learning the probability representation of the global landscape in lower dimension:

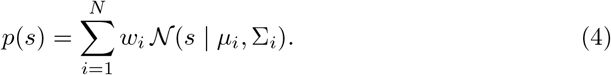

where *w*_*i*_ is the weight of the *i*-th component and *N* denotes the multivariate Gaussian probability density function for component *i* with mean *µ*_*i*_ and diagonal covariance matrix Σ_*i*_. The number of GMM components was capped at a maximum of 10 for convenience and simplification.

The residual variance not captured by the PCA components was modeled as unstructured “white noise” by calculating the 99th percentile of the absolute per-bin residual error, providing an estimate of its magnitude.

Finally, to offer depth-agnostic reconstruction, the model learns a quantitative relationship between sequencing depth (*D*) and pattern magnitude, confirming that the profile standard deviation is inversely proportional to the square root of the sequencing depth.

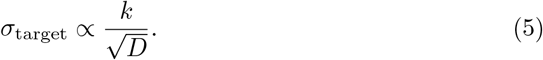

where *k* was learned from the training data by linear regression relating observed noise standard deviation to *D*^*−*1*/*2^.

During profile sampling, the GNM takes a target sequencing depth as an input and first generates a core profile by sampling from the trained PCA-GMM model. Next, several augmentations are applied to create a more diverse manifold and to allow further generalization to potential unseen batches. First, a random scaling stretch factor (*ς*) drawn from a log-normal distribution is applied to the GMM sample in latent space to modulate the intensity of the structured global patterns:

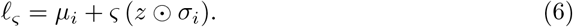

A component *i* is chosen according to the weights *w*_*i*_; *z* is a vector of random numbers drawn from a standard normal distribution, *σ*_*i*_ is the component standard-deviation vector, and ⊙ denotes element-wise multiplication.

The augmented latent vector is projected back into the high-dimensional bin space to produce the core vector:

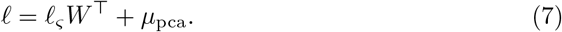

Second, an unstructured residual noise vector *ϵ* is sampled and scaled by a random factor (*υ*), also drawn from a log-normal distribution. To reproduce sharp, localized artifacts, the model also adds a synthetic spike vector *K*. The number of spikes is drawn from a Poisson distribution, and each spike has a variable width, position, and amplitude, and is shaped by a Hann window [36].

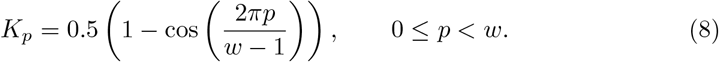

Lastly, the GNM uses the learned depth relationship to calculate a final target profile magnitude appropriate for the input depth and applies this scaling to the completed profile.

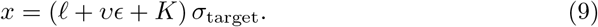

### 2.4 NPE model architecture and training

The purNPE posterior estimator was trained on simulated tumor-signal profiles augmented with GNM noise draws, without labeled cancer cfDNA samples, using a negative log-likelihood (NLL) objective. To ensure unbiased evaluation of the model’s generalization capabilities, the dataset of 250,000 bulk CNPs was split into training, validation, and testing sets based on the original 1,000 tumor evolutionary histories. This history-level split is crucial in preventing data leakage, as it guarantees that the model is validated and tested on genomic architectures derived from evolutionary histories entirely unseen during training. The final split consisted of training set 700, validation set 200, and test set 100 histories.

During training we employed an on-the-fly noise augmentation. For each clean, simulated CNP presented to the model, a new, unique noise profile was generated by sampling the GNM and added to the clean signal. This process, repeated for every sample in every epoch, exposes the model to an ever-changing manifold of sequencing noise, promoting robust learning and preventing overfitting to any specific noise pattern. In contrast, the validation set used a different approach to ensure a stable and deterministic evaluation metric. A fixed noise profile was generated and added to each of the clean validation CNPs. Using a fixed noise realization ensures that fluctuations in the validation loss are due to changes in model performance, not the stochasticity of the noise generation process, allowing for a reliable assessment of convergence.

The purNPE architecture is a conditional density estimator composed of two main components: Encoder - a deep one-dimensional Residual Network (ResNet1D) [37] acts as a feature extractor, processing the entire binned input profile and compressing it into a fixed-size summary 128-dimensional embedding. We performed a deliberate exclusion of any batch or layer normalization (e.g., BatchNorm, LayerNorm). In the context of copy number data, the absolute magnitude of the log_2_-ratio signal is directly proportional to the biological signal of interest. Normalization, although common and usually recommended for stable training regimes, would risk “squashing” or obscuring this critical magnitude information, preventing the model from distinguishing between subtle, low-amplitude alterations and high-amplitude, chaotic events. By omitting normalization, the network is forced to learn the biological significance of the signal’s raw magnitude directly, preserving the integrity of the input data.

Secondly, a density estimator - a Neural Spline Flow (NSF) [34] takes the summary embedding as a condition to model the full, joint posterior probability distribution

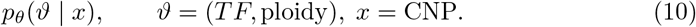

The model was trained to minimize the NLL of the true simulated parameters *ϑ* given the noisy input CNP. We used the Adam optimizer with a learning rate of 1 *×*10^*−*4^. To smooth out minor fluctuations, an exponential moving average (EMA) of the validation loss was used as the primary metric. Model selection was performed on the validation set using a composite metric that balances likelihood fit with posterior calibration and sharpness. Specifically, at each training epoch we computed:

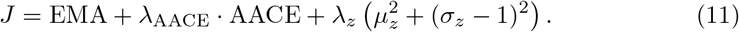

where AACE is the average absolute calibration error across coverage levels, and *µ*_*z*_ and *σ*_*z*_ are z-score diagnostics (mean and standard deviation; ideal values 0 and 1, respectively). Weights *λ*_AACE_ and *λ*_*z*_ were fixed a priori and held constant 0.2 across experiments. The best checkpoint was selected as the epoch minimizing *J* on the validation set. This procedure prioritizes likelihood while penalizing under-coverage and miscalibration. The test set was used only once for final evaluation with the selected checkpoint.

The entire training process was performed on a single NVIDIA RTX 3090 GPU. The training script was written using PyTorch and the sbi package [34] for building the estimator and posterior; exact software versions, command-line entry points, and environment files are provided with the archived code snapshot described in the Availability of data and materials section.

## 3 Results

### 3.1 A simulation-trained Bayesian framework for joint TF and ploidy estimation

The Bayesian framework is composed of two core, decoupled components: a generative forward model to produce high-fidelity training data, and an inference model trained to invert this process (Fig. 1). To generate the signal component of the training data, we created a diverse library of 250,000 biologically realistic, clean bulk CNPs. This was achieved by first generating 1,000 distinct tumor evolutionary histories using the CNAsim single-cell simulator, which models cell lineage trees under a neutral coalescent process [26]. In an attempt to capture as wide a spectrum as possible of cancer biology, we defined five distinct evolutionary archetypes ranging from genomically quiet diploid tumors to chaotic, whole genome doubled tumors with ongoing CIN [16]. This process yielded a diverse library of single-cell CNPs representing varied clonal structures and multi-scale somatic CNA patterns. To expand the diversity further, these were then computationally mixed with random combinations of clonality ratios within each evolutionary history and according to a TF drawn from a beta distribution heavily skewed towards the lower TF values (*<* 15%) to create the final clean bulk profiles (Fig. 1a).

**Fig. 1.**
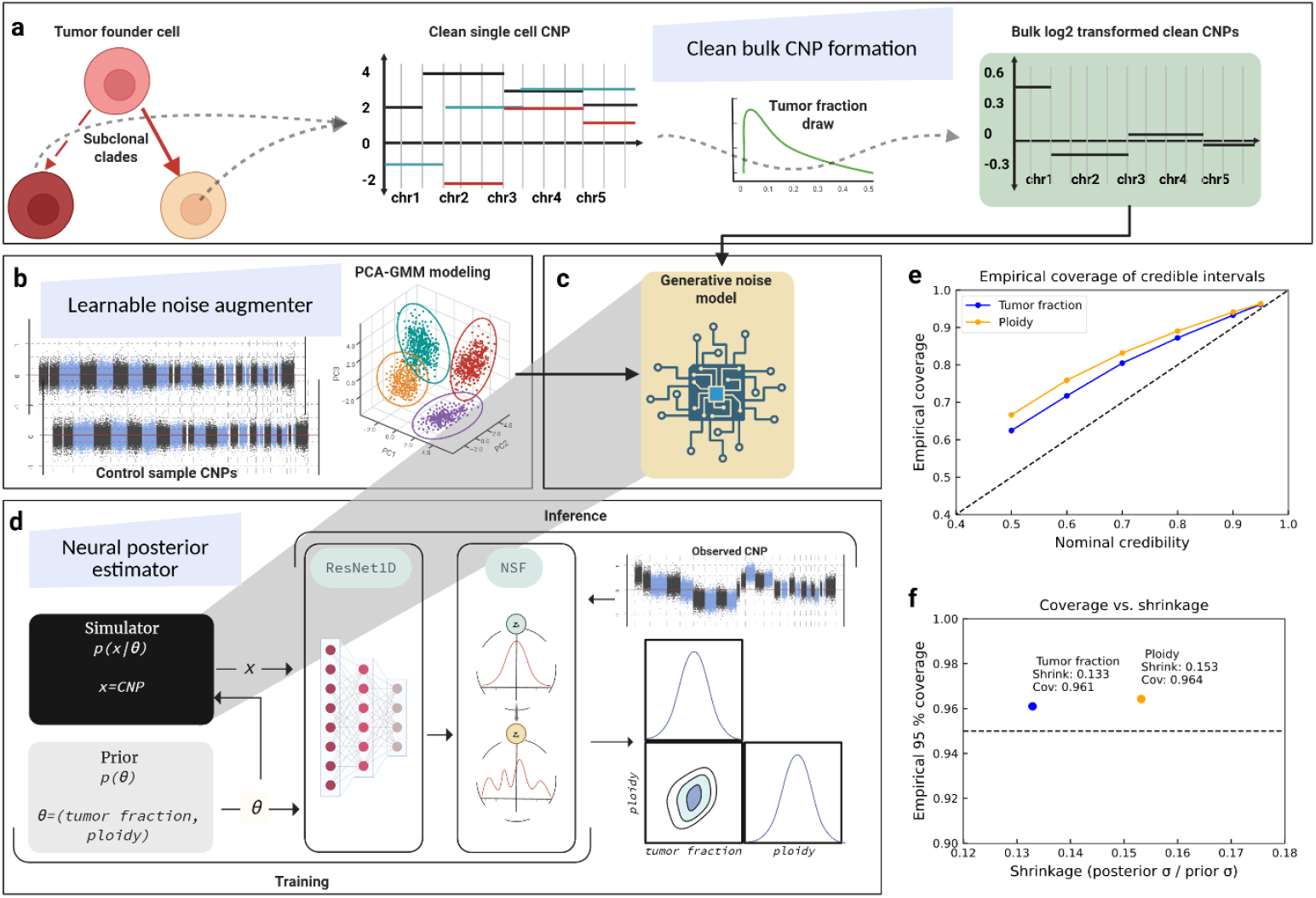
Framework schematics for the simulation-based Bayesian estimation of TF and ploidy. **a** The framework’s forward simulator generates realism-driven, clean CNPs. A tumor’s evolutionary history, including the emergence of subclones, is modelled using a coalescent-based process creating a diverse library of single-cell profiles reflecting a wide range of cancer complexities. These clean single-cell CNPs are then computationally mixed according to a TF drawn from a broad beta prior distribution, resulting in a clean bulk CNP. **b** A data-driven GNM augments the clean signal with learned sequencing artifacts. This model is a PCA-GMM trained on batches of real cancer-free control cfDNA samples, allowing it to learn and reproduce structured global sequencing patterns. **c** During training input x is the sum of the clean log_2_-transformed CNP and on-the-fly random noise augmentation. **d** The core of the framework is a NPE, which is trained to invert this process. It learns to approximate the full, joint posterior probability distribution *p*(*θ* | *x*) over the parameters *θ* = (*T F*, ploidy) given a noisy observed CNP *x*. The NPE consists of a ResNet1D encoder and a subsequent NSF, which together provide a sampling distribution over TF and ploidy. **e** The Bayesian output of the NPE is validated using simulation-based diagnostics. Empirical coverage plots for TF (blue) and ploidy (orange) show that the observed coverage of credible intervals closely tracks the nominal coverage levels. **f** Shrinkage analysis confirms the model learns substantial information from the data. The posterior standard deviation is reduced to approximately 13–15% of the prior standard deviation for TF and ploidy, respectively. Shrinkage is achieved while maintaining calibration, with over 96% of true parameters falling within the 95% credible intervals. [41]

To mitigate “sim-to-real” transfer, we developed a data-driven GNM to augment the clean bulk CNP signal with realistically learned sequencing artifacts. The model’s core is a batch-specific PCA-GMM trained on several distinct experiments, or batches, of cancer-free control cfDNA samples [3, 12, 38] (Supplementary Table S5), with the aim of learning genome wide scaffolding of the main global trends and reproduce the main sources of variance in cfDNA sequencing (Fig. 1b). Crucially, to generalize rather than overfitting to the training samples, the generative process incorporates several on-the-fly stochastic augmentations designed to create out-of-manifold examples. These augmentations include randomly scaling the magnitude of unstructured white noise, stretching the structured noise components in the latent PCA space, and adding synthetic, localized spike artifacts of variable width and amplitude. This multi-faceted approach aims to produce a noise profile that recapitulates the global properties of real sequencing data while exposing the training process to a broader noise manifold (Supplementary Fig. S1).

The core of our framework is an NPE trained without labeled cancer cfDNA samples to approximate the full, joint posterior probability distribution *p*(*TF*, ploidy| *CNP*) given a noisy, observed CNP (Fig. 1c, d). The NPE employs a deep ResNet1D to encode the entire binned read-depth profile into a fixed-size summary embedding, which then conditions a NSF [39] to model a surrogate of the posterior distribution [33] *q*_*ϕ*_(*TF*, ploidy |*CNP*) (Supplementary Table S1). This architecture provides a posterior that allows both point estimation via the median and uncertainty quantification, delivering a credible interval for each estimate.

We assessed calibration and informativeness using simulation-based diagnostics [34, 40]. Empirical coverage across credibility levels closely tracked nominal and was slightly conservative (e.g., 95% intervals covered ∼ 96% of true values; Fig. 1e). Posterior uncertainty contracted substantially relative to the prior (13–15%), demonstrating that the model extracts informative signal from complex inputs (Fig. 1f). These results indicate a statistically stable, uncertainty-aware estimator for TF and ploidy.

### 3.2 Quantification of TF and ploidy in a semi-synthetic benchmark

Because the tumor signal model was simulated and the posterior estimator was trained without labeled cancer cfDNA samples, we aimed to initially evaluate “sim-to-real” transfer at the profile level. We designed a semi-synthetic controlled spike-in experiment. This approach allows for a more direct assessment against known ground truth by combining real tumor genomic structures with an unseen sequencing-noise profile that remains constant through testing. For the noise component, we used the GC- and mappability-corrected [42] log_2_ read-depth profile from a randomly selected cancer-free control cfDNA sample from our PDAC cohort, ensuring this noise profile was entirely held out from GNM training.

To validate that the model, despite being trained on simulated data, can reliably estimate TF and ploidy from the CNPs of cancer genomes, we sourced 100 diverse, allele-specific CNPs from The Cancer Genome Atlas (GDC) [43]. We aimed to maintain a wide spectrum of genomic instability by selecting a spectrum of profiles with Percent Genome Altered (PGA) from genomically quiet (minimum PGA = 0.09) to highly chaotic (maximum PGA = 0.99) genomes (Supplementary Fig. S2; Supplementary Table S2). Each of these 100 real cancer CNPs was computationally injected into the control profile’s log_2_ read-depth space at 30 discrete TF levels, ranging from 0.5% to 15% in 0.5% intervals.

We ran inference on all 3,000 resulting augmented genomes and compared its performance to ichorCNA [7], the leading TF caller based on binned read-depth analysis and as such the optimal contender. We ran ichorCNA in two distinct modes: first, a generic execution using its default provided Panel of Normals (PoN) and the minimal 500-kb bins (ichorCNA 500-kb), this mode is regarded by us as the most appropriate competition, under these settings both methods aim to infer low-TF without a sample-specific matched control during inference. Additionally, an optimally configured execution (ichorCNA) that utilized a cohort-matched PoN built from the remaining 11 control samples and a higher resolution of 50-kb bins. This mode leverages reduction of the batch specific sequencing artifacts via standardized methods, a process that can be substantial in vast scenarios. Both modes were executed using wide parameters search which proved optimal for accurate low-TF estimates [44] (Extended methods).

The results of this experiment are visualized with two representative examples in Fig. 2. We selected a genomically quiet mesothelioma profile (PGA = 0.13, Fig. 2a) and a chaotic, WGD testicular cancer profile (PGA = 0.99, Fig. 2b). Even at a 7% TF, the underlying CNA signal is subtle and difficult to discern from the baseline noise in the low-PGA case, whereas it is prominent in the high-PGA sample. Head-to-head comparisons across the full TF range (Fig. 2c, d) demonstrate that our model’s median estimates (blue line) closely track the ground-truth (dashed line) for both TF and ploidy. Notably, the 95% credible intervals (shaded blue area) accurately reflect the underlying uncertainty, appearing wider for the challenging low-PGA profile and narrower for the high-signal, high-PGA profile. In contrast, both ichorCNA modes show underestimation of TF and volatility in ploidy estimates, particularly at low signal levels.

**Fig. 2.**
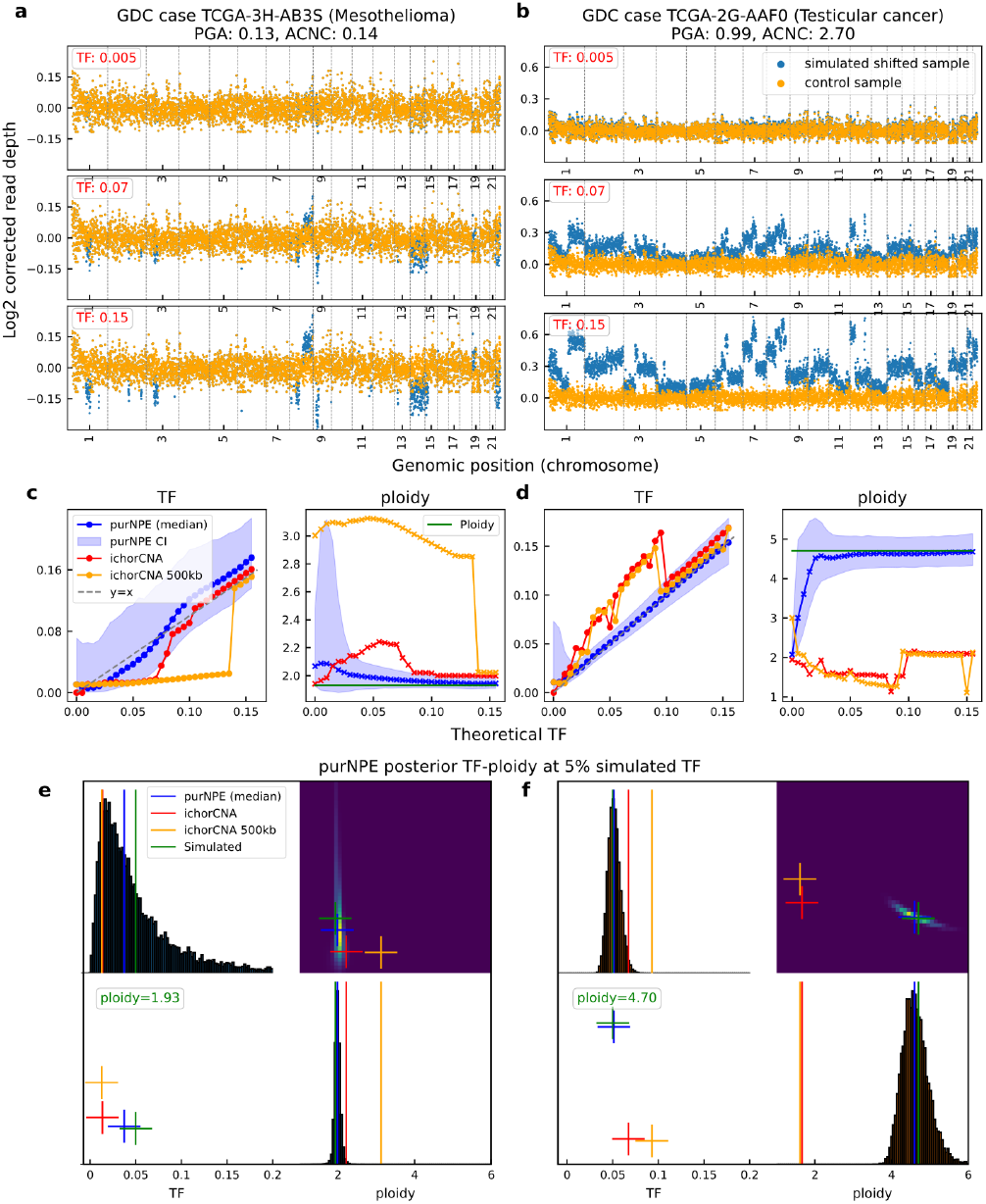
TF and ploidy estimation in a semi-synthetic cohort using real cancer genomes and unseen noise pattern. **a, b** Representative genome-wide copy number profiles from the in silico spike-in experiment. A genomically quiet mesothelioma profile (low PGA, **a**) and a chaotic testicular cancer profile (high PGA, **b**) were computationally mixed into a real, unseen cancer-free control cfDNA sample at TFs of 0.5%, 7%, and 15%. Blue dots represent the log_2_-transformed read depth of the mixed sample, while orange dots show the baseline noise of the control sample. Even at 7% TF, the CNA signal is subtle in the low-PGA case but prominent in the high-PGA case. **c, d** Head-to-head comparison of estimation performance across a range of TFs (0–15%) for the low-PGA (**c**) and high-PGA (**d**) cases. PurNPE’s median estimate (blue line) closely tracks the ground truth y=x line for both TF and ploidy. The 95% CI (shaded blue) reflects the higher uncertainty in the low-PGA setting. In contrast, ichorCNA (red and orange lines, representing two different execution modes) show underestimation of TF and volatility in ploidy estimates, particularly at low signal levels. **e, f** Full Bayesian posterior distributions from purNPE for a 5% simulated TF. The plots show the joint posterior (heatmap) and the marginal distributions (histograms) for TF and ploidy for the low-PGA (**e**) and high-PGA (**f**) cases. PurNPE’s posterior is centered on the true simulated values (green lines), providing an uncertainty-aware output. Point estimates from ichorCNA competitor methods are given in red and orange lines.

To highlight the advantage of a full Bayesian framework over point estimates, we visualized the complete joint posterior distribution from the model for a 5% TF spike-in (Fig. 2e, f). The posterior is tightly centered on the true simulated values (green lines), providing an uncertainty-aware output.

Next, we performed statistical analysis of the semi-synthetic spike-in cohort of 3,000 simulated genomes (Fig. 3). We first stratified the Mean Absolute Error (MAE) of TF prediction by both ground-truth TF and cancer genome complexity, proxied by the PGA values (Fig. 3a). Our model maintained low prediction error (*<* 3% MAE) across conditions while both ichorCNA modes exhibited variable error patterns; notably, in high-PGA genomes both modes showed systematic reduction in estimating TF of genomically chaotic tumors.

**Fig. 3.**
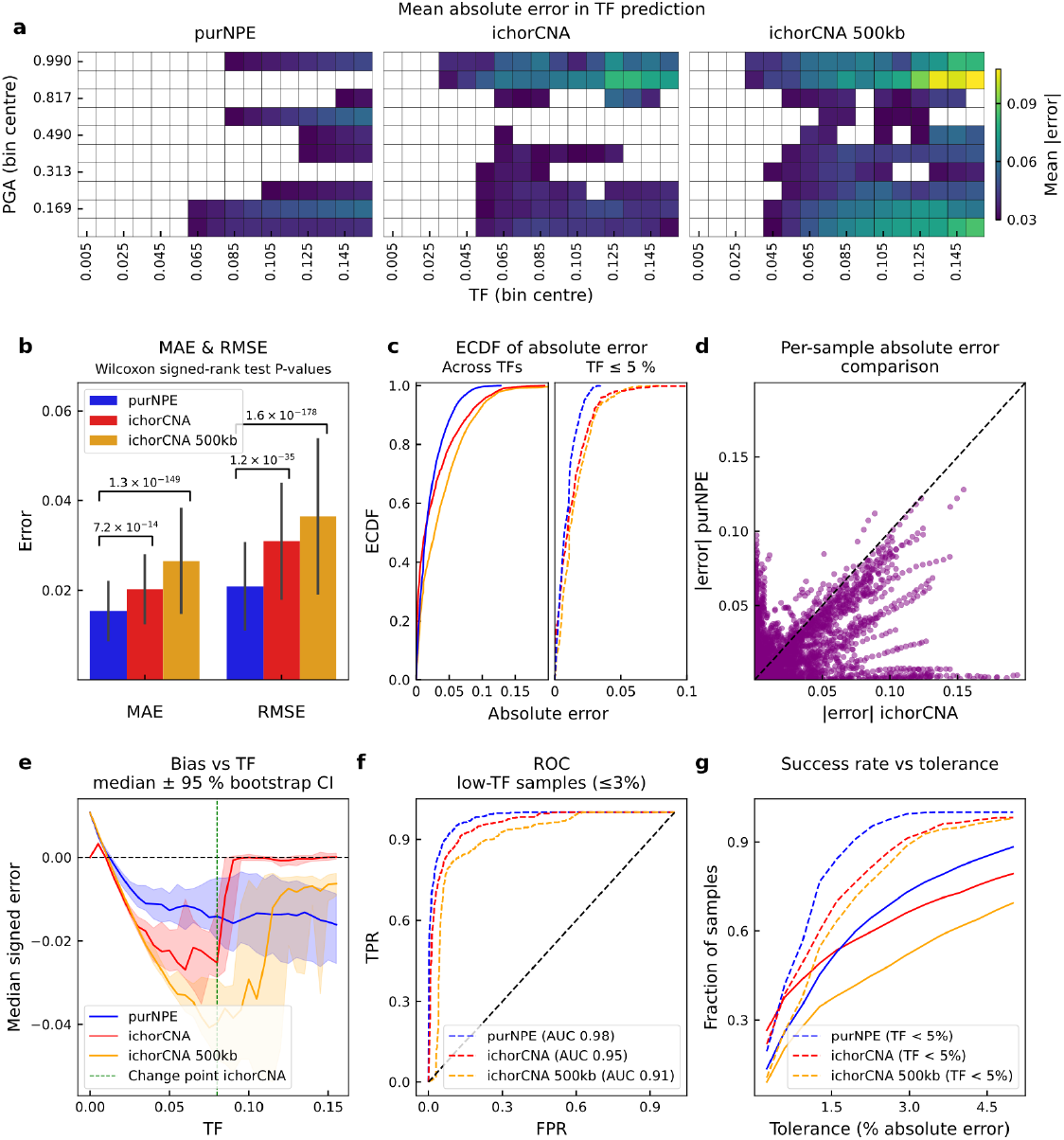
Cumulative statistical validation on the semi-synthetic cohort for the assessment of accuracy, bias estimates, and reliability. **a** MAE of TF prediction, stratified by ground-truth TF (x-axis) and cancer genome complexity (PGA; y-axis). Each cell represents the average error across hundreds of simulations for that combination. Bins with a mean error *<* 3% are masked (white) for clarity. PurNPE maintains low error (dark purple) across a wide range of conditions, whereas ichorCNA and ichorCNA 500-kb show a more pronounced error (green/yellow), particularly for high-PGA genomes. **b** Aggregate MAE and RMSE across all simulations. PurNPE achieves lower error (nominal Wilcoxon signed-rank test *P* values shown). Error bars represent the 95% confidence interval of the mean. **c** ECDF of absolute errors. The blue curve (purNPE) is consistently to the left, indicating a higher proportion of predictions with smaller errors in the clinically relevant low-TF (*≤* 5%) regime (dashed lines). **d** Per-sample comparison of absolute errors between purNPE and ichorCNA. The majority of points lie below the y=x line indicating that for most individual simulations, purNPE’s predictions are more accurate. **e** Median signed error as a function of TF, with 95% bootstrap confidence intervals. PurNPE demonstrates a median error centered around - 0.01 across TFs. In contrast, both ichorCNA modes exhibit an underestimation at TFs below ∼8%. **f** ROC curve for the task of classifying low-fraction samples (*≤* 3% TF) from cancer-free controls (0%). PurNPE achieves an AUC of 0.98. **g** Success rate vs. tolerance, showing the fraction of samples with a prediction error less than or equal to a given tolerance. PurNPE’s success rate is consistently higher.

The aggregate results demonstrated lower overall MAE (2.2%) for the Bayesian model compared to ichorCNA (2.8%) and ichorCNA 500-kb (3.8%) (Wilcoxon signed-rank test between purNPE and ichorCNA modes, *P* = 7.2 ×10^*−*14^ and *P* = 1.3× 10^*−*149^, respectively, Fig. 3b). Because multiple semi-synthetic profiles share source tumor genomes, dilution levels, and a fixed background-noise profile, these inferential *P* values should be interpreted descriptively; we therefore emphasize effect sizes and empirical error distributions. To assess overall accuracy, the Empirical Cumulative Distribution Function (ECDF) of absolute errors (Fig. 3c) was calculated; here the curve is consistently shifted to the left, indicating a greater proportion of predictions with minimal error. In the clinically relevant low-TF ( ≤ 5%) regime (dashed lines), our model’s curve rises more steeply than those of its competitors. A per-sample comparison between purNPE and ichorCNA shows that on this specific semi-synthetic benchmark the majority of data points fall below the identity line, indicating that the Bayesian framework was more accurate on an individual basis (Fig. 3d).

We found that purNPE maintains a roughly consistent bias of -0.01 median signed error across the entire TF range (Fig. 3e). Conversely, both ichorCNA modes displayed a systematic underestimation bias in the low-TF regions and recovering with increasing TF values (*>* 8%) and approaching zero. Furthermore, we calculated classification ability of the cohort by distinguishing low-TF samples (≤ 3%) from cancer-free controls (TF = 0%). Here, purNPE achieved an Area Under the Curve (AUC) of 0.98, compared to 0.95 and 0.91 by ichorCNA and ichorCNA 500-kb, respectively (Fig. 3f). Furthermore, purNPE maintains consistency with higher success rate, the fraction of samples predicted within a given tolerance, most notably in the TF *<* 5% subset (Fig. 3g).

Finally, we checked the statistical validity of the Bayesian framework in this controlled semi-synthetic setting. An uncertainty calibration plot demonstrated that the model’s reported 95% credible interval contains the true parameter value approximately 95% of the time (Supplementary Fig. S3a). The Bayesian framework displays a symmetrical distribution of estimates around the ground-truth *y* = *x* line in the challenging low-TF (≤ 5%) regime (Supplementary Fig. S3b). Together, these results provide controlled evidence regarding the transferability of the simulated CNPs to real profiles while benchmarking the performance of purNPE compared to the closest existing method.

### 3.3 Accuracy and uncertainty estimates against an orthogonal MAF proxy in a public pan-cancer cohort

To assess the generalizability and performance of the model in a real-world setting, we analyzed a diverse public pan-cancer clinical cohort of 106 cfDNA samples from the study by Mouliere et al. [12] (Supplementary Table S6). For this cohort, the maximum MAF, determined by either digital PCR (dPCR) or hybrid-capture deep sequencing, serves as an orthogonal biological proxy for TF. We benchmarked our approach against the two previously described ichorCNA modes, with a dedicated PoN built for the ichorCNA variant from samples provided as cancer-free in the study. We included two additional methods: t-MAD, a metric from the original Mouliere et al. study calculated as the “trimmed median absolute deviation from copy number neutrality” from shallow WGS, and FRAGLE [11], a model that predicts TF by analyzing the density distribution of cfDNA fragment lengths. Lastly, we excluded all cancer-free samples from the Mouliere et al. study from the GNM during training to avoid leakage of similarly behaving profiles and to preserve assessment of inference without a cohort-specific PoN.

When applied to this cohort, our model’s TF estimates demonstrated a statistically significant correlation with the orthogonal MAF proxy (Pearson’s *r* = 0.734, *P* = 3.6 ×10^*−*19^) (Fig. 4a). As previously mentioned, the unique advantage of the Bayesian framework is the ability to provide a full posterior probability distribution that can be summarized by a credible interval. When we evaluated this output, we found that the orthogonal MAF proxy was contained within the model’s 95% credible interval in 75.7% of cases with MAF ≤ 15% (71.7% overall) (Fig. 4b). This result should be interpreted as agreement with an imperfect biological proxy rather than definitive posterior calibration against true TF. Furthermore, credible intervals were wider for samples with low MAF as the signal of aneuploidy diminished relative to the overall noise, indicating increased model uncertainty about the TF. This contrasts with alternative non-Bayesian methods that provide a single point estimate without context of the potential error.

**Fig. 4.**
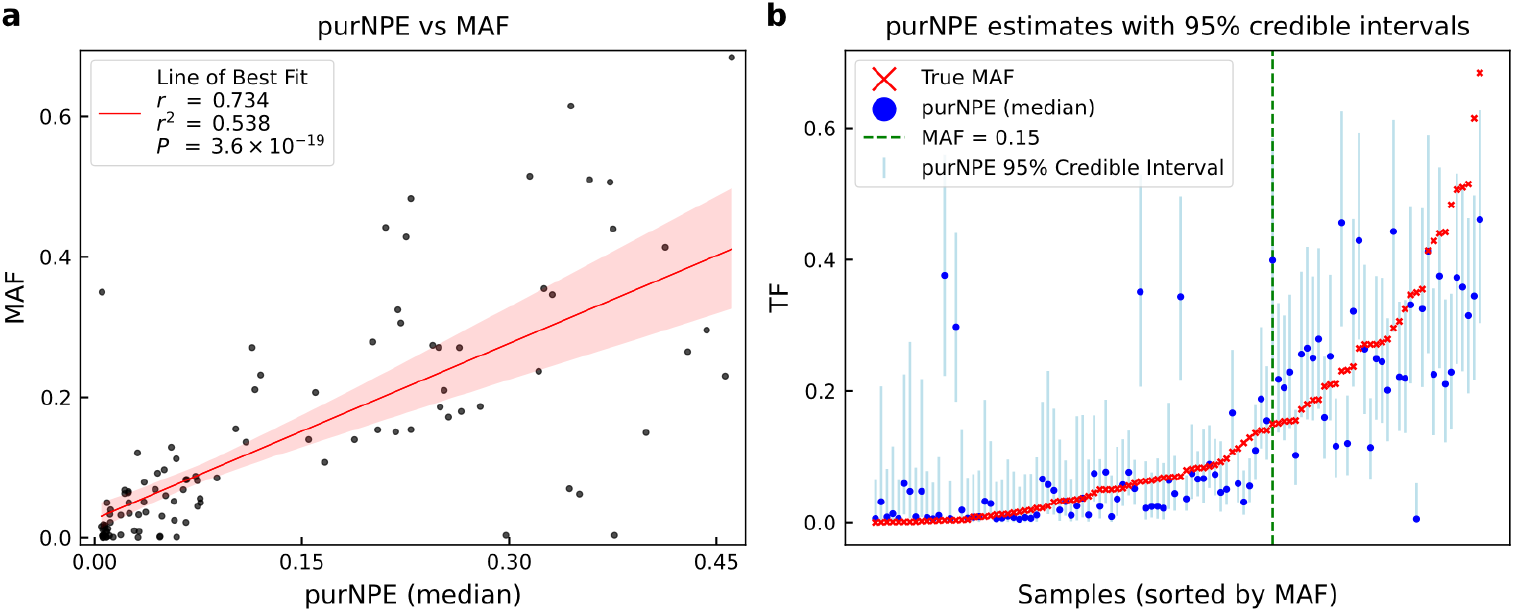
Performance on a real-world pan-cancer clinical cohort. **a** Correlation between TF estimates from purNPE and an orthogonal MAF proxy. The analysis was performed on a cohort of 106 clinical cfDNA samples from a pan-cancer study by Mouliere et al. PurNPE’s estimates show a statistically significant correlation with MAF (Pearson’s *r* = 0.734, *P* = 3.6 ×10^*−*19^), demonstrating agreement across a diverse set of real-world cancer types. The solid red line indicates the line of best fit, with the shaded area representing the 95% confidence interval of the regression. **b** Per-sample visualization of purNPE’s Bayesian output for the Mouliere et al. cohort. For each sample (sorted by increasing MAF), the MAF proxy (red “x’) is shown alongside purNPE’s median estimate (blue dot) and its 95% credible interval (light blue bar). The credible intervals reflect higher uncertainty for low-MAF samples. Across samples with MAF ≤ 15% (dashed green line), the MAF proxy is contained within purNPE’s 95% credible interval in 75.7% of cases; this is an interval-agreement analysis against a proxy, not definitive posterior calibration against true TF.

In comparison to the competitors, purNPE achieved favorable point metrics (Table 1). It demonstrated the highest Pearson correlation coefficient and the lowest MAE (0.066). While the improvement in absolute error over the optimally configured ichorCNA did not reach statistical significance (Wilcoxon signed-rank test on absolute errors, *P* = 0.559), purNPE achieved these metrics without requiring a matched control sample or cohort-specific PoN at inference and outperformed the more comparable ichorCNA 500-kb mode (*P* = 6.51*×* 10^*−*7^). The full correlation plots for all competitor methods are given in Supplementary Fig. S4. This includes an analysis restricted to the “high signal” subset as defined in the original Mouliere et al. study, where the Bayesian method still demonstrates the highest correlation. Furthermore, a visual performance assessment of the model trained on clean CNPs only under an otherwise identical training regime is displayed. This showcases the crucial influence of the GNM in calibrating the model to sequencing depth and genome-wide trends.

**Table 1.**
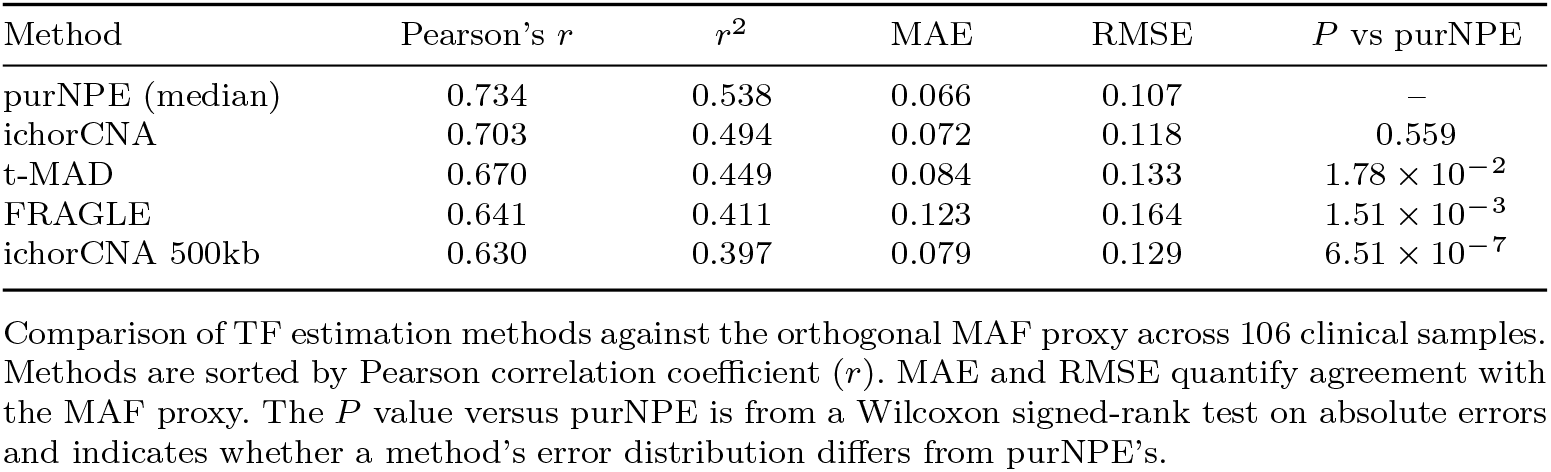
Performance comparison on a clinical cohort with orthogonal MAF proxy.

| Method | Pearson’s $r$ | $r^2$ | MAE | RMSE | $P$ vs purNPE |
| --- | --- | --- | --- | --- | --- |
| purNPE (median) | 0.734 | 0.538 | 0.066 | 0.107 | – |
| ichorCNA | 0.703 | 0.494 | 0.072 | 0.118 | 0.559 |
| t-MAD | 0.670 | 0.449 | 0.084 | 0.133 | $1.78 \times 10^{-2}$ |
| FRAGLE | 0.641 | 0.411 | 0.123 | 0.164 | $1.51 \times 10^{-3}$ |
| ichorCNA 500kb | 0.630 | 0.397 | 0.079 | 0.129 | $6.51 \times 10^{-7}$ |

Comparison of TF estimation methods against the orthogonal MAF proxy across 106 clinical samples. Methods are sorted by Pearson correlation coefficient ( $r$ ). MAE and RMSE quantify agreement with the MAF proxy. The $P$ value versus purNPE is from a Wilcoxon signed-rank test on absolute errors and indicates whether a method’s error distribution differs from purNPE’s.

### 3.4 Concordance assessment on independent clinical data

To further assess the generalizability of the method, we evaluated performance on two independent clinical datasets: the PDAC cohort (n = 9 cases, n = 12 controls, Supplementary Table S7) and a breast cancer cohort (n = 9 cases, n = 9 controls, Supplementary Table S8). These cohorts represent a real-world data scenario, with each library prepared and sequenced at different facilities using distinct protocols (Extended methods). We compared our model with the above-mentioned competitors. For the optimally configured ichorCNA variant, we constructed a dataset-specific PoN from the matched control samples in each dataset, consequently reducing the number of samples used and inferring TF on the cancer samples only. The PDAC dataset was of mixed sequencing depth, and in the attempt to meet FRAGLE and t-MAD guidelines for shallow WGS, these samples were randomly downsampled to 10 million reads (Supplementary Table S4) while testing the two methods.

purNPE demonstrated high concordance and a linear correlation with ichorCNA (Pearson’s *r* = 0.81) and ichorCNA 500-kb (Pearson’s *r* = 0.84) (Table 2). This agreement was further quantified by a low MAE of 0.6–0.7% between the methods. A Bland-Altman analysis confirmed this consistency, revealing a low positive bias (+0.5% to +0.6%) and tight 95% Limits of Agreement (LoA) of approximately *±*1% (Fig. 5a).

In contrast to the agreement with the ichorCNA variants, purNPE showed lack of correlation with t-MAD and FRAGLE (Pearson’s *r* = −0.31 and *r* = −0.04, respectively). This systematic disagreement translated into large MAEs (9.3% and *−*4.7%) (Table 2, Supplementary Fig. S5). This pattern of disagreement is consistent with that observed between the ichorCNA variants and the remaining methods as well, signaling the potential variability between different cfDNA or cancer features and their summary. Collectively, these results indicate that purNPE shows comparable behavior to ichorCNA, a widely used read-depth estimator, on independent clinical datasets from diverse technical backgrounds.

**Table 2.** Inter-method agreement and bias analysis on two independent clinical cohorts.

| Method compared to purNPE | Pearson’s $r$ | Bias | LoA (lower, upper) | MAE | RMSE |
| --- | --- | --- | --- | --- | --- |
| ichorCNA 500kb | 0.836 | +0.005 | (0.001, 0.010) | 0.005 | 0.006 |
| ichorCNA | 0.809 | +0.006 | (−0.001, 0.012) | 0.006 | 0.007 |
| FRAGLE | −0.040 | +0.044 | (−0.052, 0.140) | 0.047 | 0.065 |
| t-MAD | −0.312 | +0.093 | (−0.031, 0.217) | 0.093 | 0.112 |

Summary of pairwise comparisons between purNPE, used as the reference method, and competitor methods across two clinical cohorts (PDAC and breast cancer). For the ichorCNA mode, a PoN was constructed from the control samples within each cohort, reducing the number of available samples for comparison. For all other modes, the full cohort was used. Bias and 95% LoA are derived from Bland–Altman analysis. MAE quantifies the average magnitude of the difference.

**Fig. 5.**
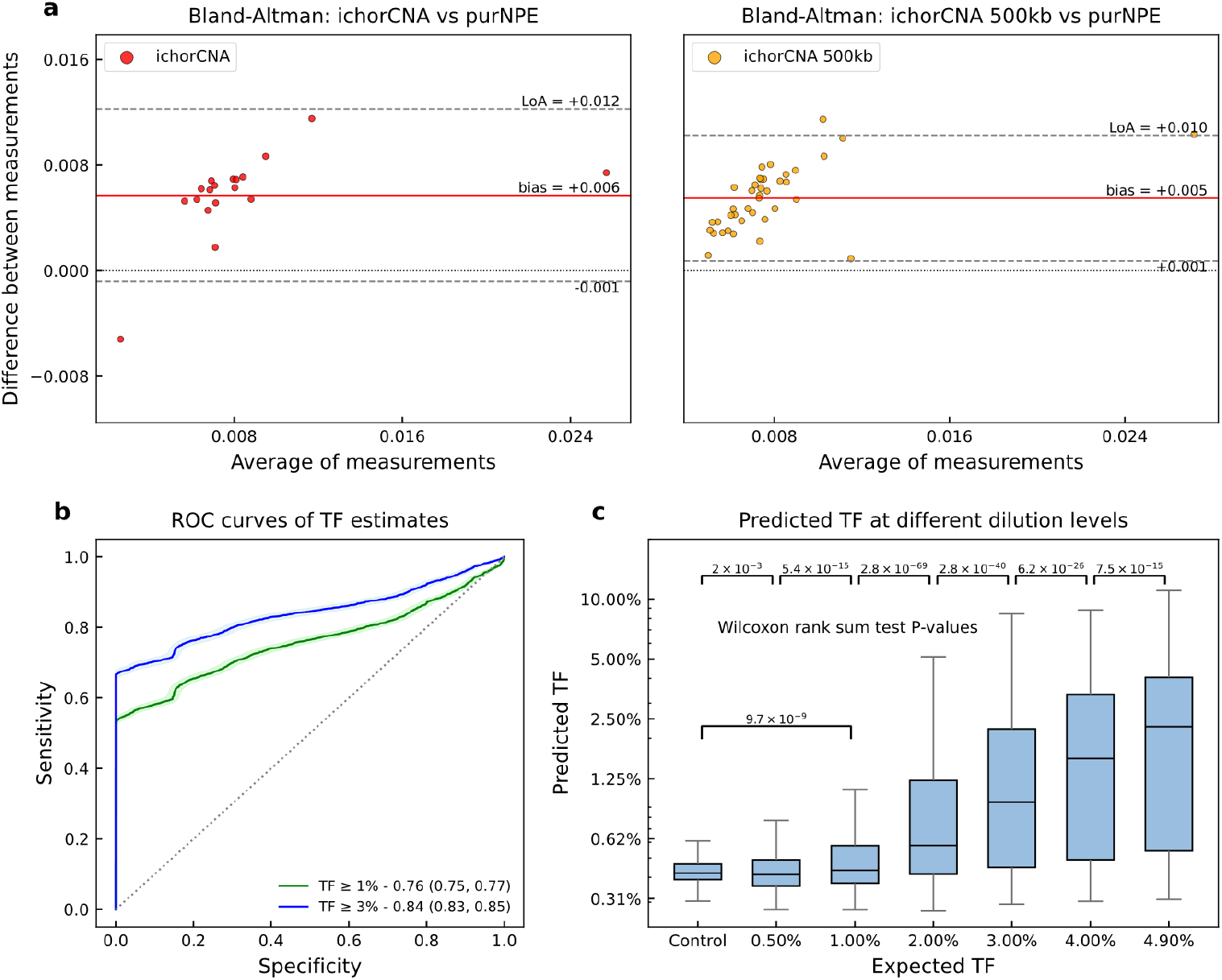
Analytical sensitivity in the low-TF regime and concordance analysis with existing methods on independent clinical datasets. **a** Bland-Altman plots comparing TF estimates from purNPE and two modes of ichorCNA on two independent clinical cohorts (PDAC and breast cancer). The low bias (+0.005 to +0.006) and tight 95% LoA demonstrate concordance between purNPE’s estimates and those from the ichorCNA variants. **b** ROC curves for classifying control samples versus samples with low TF. The classification task was performed on the in silico cohort generated by spiking the GDC profiles into the PDAC and breast cancer cancer-free control samples. PurNPE achieves an AUC of 0.76 (95% CI, 0.75–0.77) for detecting TF ≥1% and 0.84 (95% CI, 0.83–0.85). **c** Predicted TF at different expected dilution levels from the same in silico cohort. The distribution of predicted TF shows a dose-response and is statistically separable from the baseline control group (nominal Wilcoxon rank-sum test *P* values shown). The difference between the control and 1% groups (*P* = 9.7 ×10^*−*9^) suggests analytical sensitivity near 1% under the evaluated semi-synthetic conditions. Boxplots represent the median and interquartile range.

### 3.5 Semi-synthetic low-fraction analyses suggest analytical sensitivity near 1% TF

A critical attribute for any TF estimator is its sensitivity in the analytically challenging low-TF regime, which is relevant for applications involving low disease burden. We assessed purNPE’s sensitivity using an in silico experiment generated by spiking the 100 GDC profiles into the cancer-free control cfDNA samples from the PDAC and breast cancer datasets, resulting in 21 distinct control backgrounds. Consistent with the leakage-control strategy described above, the model was trained while excluding the matching cancer-free samples of the PDAC and breast cancer cohorts from GNM augmentation.

First, we evaluated the model’s performance as a classifier for detecting low-TF samples from controls. PurNPE demonstrated moderate classification capability, achieving an AUC of 0.76 (95% CI, 0.75–0.77) for identifying samples with TF≥ 1% and a higher AUC of 0.84 (95% CI, 0.83–0.85) for those with TF≥ 3% (Fig. 5b). The ability to separate low-TF groups from the baseline controls was statistically significant (Mann–Whitney *U, P* = 1.0×10^*−*293^); because these spike-ins reuse source tumor profiles and control backgrounds, the *P* value should be interpreted descriptively.

To estimate low-fraction analytical sensitivity, we analyzed the model’s performance on a dilution series ranging from 0.5% to 4.9% TF. TF estimates exhibited a clear dose-response, with the median prediction increasing monotonically with each successive dilution level (Fig. 5c). The distribution of purNPE’s estimates for the 1% dilution group was separable from the baseline control group (Wilcoxon rank-sum test, *P* = 9.7 *×*10^*−*9^). These results suggest analytical sensitivity near 1% TF under the evaluated semi-synthetic conditions, but do not constitute prospective clinical LoD validation.

Analytical sensitivity was also estimated by determining the Limit of Blank (LoB) and LoD according to CLSI EP17-A2 guidelines [45] (Extended methods). Based on 900 blank and 900 low-level semi-synthetic replicates, the LoB was 0.56% TF, and the LoD estimate was 1.16% TF. This additional semi-synthetic sensitivity analysis supports the conclusion that the model can separate approximately 1% TF spike-ins from blanks under the evaluated conditions.

## 4 Discussion

This study evaluates whether simulated tumor copy-number signals, paired with learned cfDNA sequencing-noise patterns, can support tumor-fraction estimation in a data-scarce setting. SBI has been demonstrated to be effective across fields of science where sampling large quantities of simulated data from a realistic-enough prior can be useful in training deep neural Bayesian frameworks [46–49]. While DL methods have been previously applied to cfDNA analysis [11, 50–52], these methods, for the most part, utilize supervised learning regimes and as a result are limited to the manifold spanned by the curated training set. Moreover, deep neural network training is data intensive, and securing a rich set of diverse, accurately labeled samples is a constant challenge in oncology and healthcare sciences at large. Simulations, assuming sufficient diversity and fidelity are attainable, can reduce this dependence on labeled cancer cfDNA cohorts.

The statistical distribution of CNAs and the progression of clonal evolution have been investigated for many years, resulting in mathematical evolutionary models and statistical summaries of plausible aberrational genomic landscapes. Under the assumption that genome-wide copy-number dispersion can be treated as a pattern-recognition problem, neural networks are natural candidates for learning aneuploidy patterns from CNPs. We tested this assumption by simulating not only the clean signal of tumor aneuploidy but also global patterns characteristic of whole-genome cfDNA sequencing, through modeling major sources of variance across sequencing chemistries and platforms and stochastically augmenting global and local artifacts. This study’s central objective was to assess whether this simulation-plus-noise strategy transfers to real or semi-synthetic cfDNA profiles.

A fundamental innovation of purNPE is its shift away from the traditional paradigm of per-sample statistical fitting. Methods like ichorCNA, ASCAT, Battenberg and others, while powerful, are computationally intensive procedures that treat each sample in isolation. In contrast, purNPE is expensive to train relative to these methods, but once trained it leverages amortized inference, allowing it to make predictions in a fraction of a second with minimal computational resources while still delivering an expressive posterior. For comparison, the parameter grid search performed in the low-TF protocol of ichorCNA can take up to 3 minutes on similar hardware. This efficiency is a direct result of the model having learned recurrent statistical patterns of aneuploidy across thousands of simulated histories. This pattern-recognition approach is strengthened further by the data-driven GNM, which acts as a bridge for “sim-to-real” transfer by augmenting clean tumor signals with realistic sequencing artifacts. Under the evaluated conditions, purNPE achieved performance comparable to ichorCNA, including comparisons where ichorCNA was run in its cohort-matched configuration.

The advantage of the Bayesian framework is its ability to provide not only a point estimate for TF and ploidy but also a calibrated credible interval. This is a feature absent from many tools that either aim to find a set of parameters that maximizes a likelihood objective, in the case of ichorCNA, or the result of a deterministic calculation, such as in the case of t-MAD. In the low-TF space the signal-to-noise ratio is low, making honest quantification of uncertainty important for downstream interpretation. We contend that the wide credible intervals reported by purNPE for very low-TF, or extreme PGA, are a critical feature of a well-behaved Bayesian system reflecting its low confidence in the estimate. The semi-synthetic sensitivity analyses suggest that purNPE can separate approximately 1% TF spike-ins from blanks under the evaluated conditions; however, this should not be interpreted as prospective clinical LoD validation.

In the current implementation we made a deliberate design choice to exclude sub-clonal fraction estimation, even though it is a popular quantity to infer in related tools. Our initial experiments indicated that attempting to infer subclonality in the low-TF regime introduced unreliability into the primary TF and ploidy estimates. We therefore prioritized robustness and accuracy for the low-TF space, while recognizing that future versions could also aim to provide reliable estimates for that quantity. More generally, the purNPE framework is highly extensible. Although the posterior estimator in this version was trained without labeled cancer cfDNA samples, future iterations could be fine-tuned on curated datasets of real tumor profiles to enhance performance, adapt to domain-specific nuances, and expose the model to CNP patterns not captured by the simulator. Another solution could introduce a mixture-of-experts architecture to better accommodate different sequencing depths, TF regimes, or sequencing chemistries. The model architecture can also be modified; we experimented with several encoder architectures, including Transformer-based models and dilated convolutions, but found that our ResNet1D implementation offered the best combination of performance and efficiency. A promising next step could be an encoder-decoder architecture, where a reconstruction loss on the embedding enforces learning of a more informative CNP summary. Similarly, while NSF gave the leading results for the density estimator, other estimators or likelihood-free learners can be explored. Lastly, binned read-depth analysis constitutes only one vertical in cfDNA analysis; merging multiple feature sets from fragmentomics and epigenetic markers might improve robustness. A multi-modal approach is therefore an additional future direction for purNPE.

Even though purNPE demonstrated promising results in our evaluations, we recognize meaningful limitations. First, performance is intrinsically linked to the fidelity and realism of the simulations. We chose CNAsim due to its tumor-evolution engine, but this statistical model may miss forms of complex genomic instability, such as chromothripsis or breakage-fusion-bridge cycles, that are observed in real tumors. Our parameter randomization within the simulator space may also miss relevant evolutionary scenarios. Second, the GNM is limited by the span of sequencing chemistries and regimes represented in the cancer-free control profiles. Although dimensionality reduction and randomized augmentation generate out-of-manifold noise patterns, the model remains constrained by the patterns observed in the training controls. Third, the real-data evaluations are not equivalent to prospective clinical validation: MAF is an imperfect proxy for TF, the independent clinical cohorts are small, and the low-fraction LoD analyses are semi-synthetic. We therefore view the clinical and concordance results as preliminary and consider validation in larger, prospectively designed cohorts with orthogonal ground-truth measurements essential before clinical deployment.

## 5 Conclusions

purNPE provides a reproducible simulation-based Bayesian framework for uncertainty-aware TF and ploidy estimation from cfDNA CNPs. By combining tumor CNP simulation, learned cfDNA noise augmentation, and NPE, the method reduces dependence on large labeled cancer cfDNA training cohorts and does not require a matched control or cohort-specific PoN at inference. The current evidence supports purNPE as a methodological alternative to existing read-depth estimators under the evaluated conditions, while larger datasets with stronger orthogonal ground truth and prospective validation are needed to determine its translational utility.

## Declarations

### Ethics approval and consent to participate

This study was conducted in accordance with the ethical standards of the institutional research committees. The study was approved by the Helsinki Committee of the Sheba Medical Center (approval number 9534-22-SMC) and the Helsinki Committee of the Tel Aviv Sourasky Medical Center (approval number 0569-16-TLV). Informed consent was obtained from all participants prior to sample collection.

### Consent for publication

Not applicable.

### Availability of data and materials

The raw sequencing data generated in this study have been deposited in the European Genome-phenome Archive (EGA) under study accession number EGAS50000001620. Control samples used for training the GNM are available in the EGA under accession IDs EGAD00001005339, EGAD00001006237, and EGAD00001006132. Pretrained model weights, simulated training and validation data, processed read counts needed to reproduce model training, and a snapshot version of the tool are available from Zenodo [53]. The purNPE software is available from the project repository [54].

### Authors’ contributions

H.V. designed, planned and executed the study and drafted the manuscript. M.R.-G. and T.R. assisted in sample acquisition. M. Grad and R.S. performed the biological experiments. A.D. assisted in sample acquisition. M. Gorfine contributed statistical guidance. N.S. supervised the study. All authors reviewed the manuscript and read and approved the final version.

## Acknowledgements

We thank members of the Shomron laboratory for handling the biological processing and DNA extraction along with valuable support in data acquisition and technical consultation. We thank Dr. Moni Shahar from the AI and Data Science Center, Tel Aviv University, Israel, for valuable insights during the ideation process. We thank the TLV BioBank, Tel Aviv Sourasky Medical Center, Tel Aviv, Israel, for providing the breast cancer and control patients cohort.

