## Supplemental Materials and Methods for "Simulation-based Bayesian deep learning enables uncertainty-aware tumor fraction estimation in cell-free DNA"

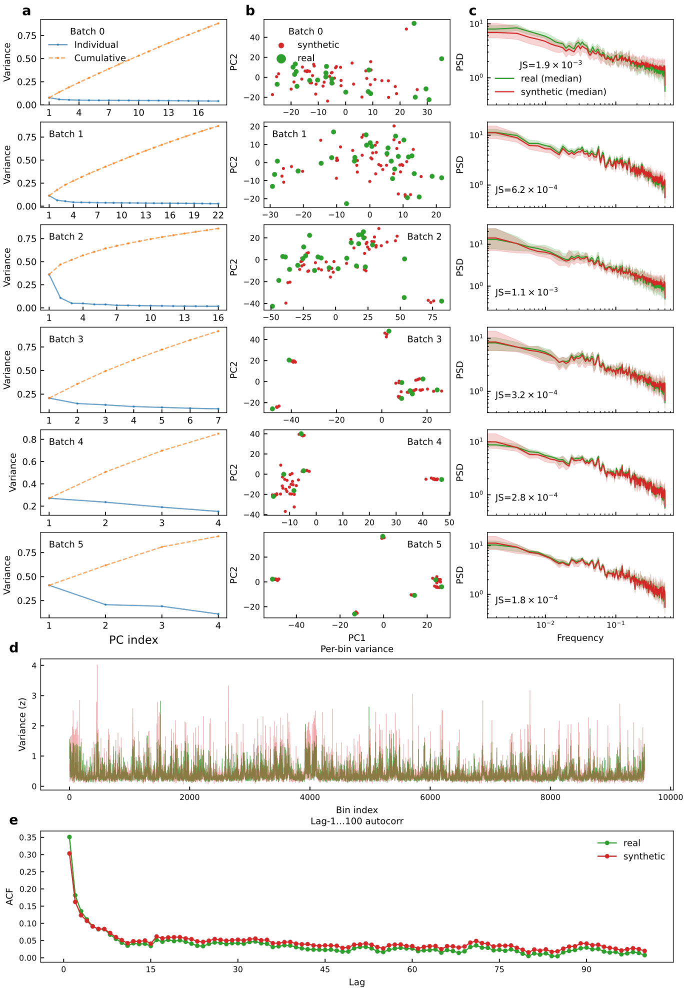


**Fig. S1** **Development and multi-scale validation of the data-driven GNM.** The GNM is trained on cancer-free control cfDNA read-depth profiles (GC- and mappability-corrected $\log_2$ ratios) and conditioned on sequencing batch. **a** Scree plots of PCA performed separately for each of the six batches used in the study show that a small number of principal components explain most of the variance, indicating low-dimensional global structure. **b** For each batch, control samples and synthetic profiles drawn from the GNM with matched read depth are projected onto the first two PCs. The overlap indicates that dominant batch structures are reproduced. **c** Batch-wise median power spectral density (Welch) with 5–95% envelopes for real and synthetic profiles. Agreement is quantified by Jensen–Shannon (JS) divergence between the median PSDs. **d** Genome-wide per-bin variance, on a $z$-score scale, aggregated across all batches for real and synthetic profiles, demonstrating that the model reproduces position-specific variability. **e** Mean unbiased autocorrelation function for lags 1–100 across samples, showing agreement of short- and long-range correlations between real and synthetic data. Together, these visualizations show that the GNM reproduces the global trend structure of cfDNA sequencing.

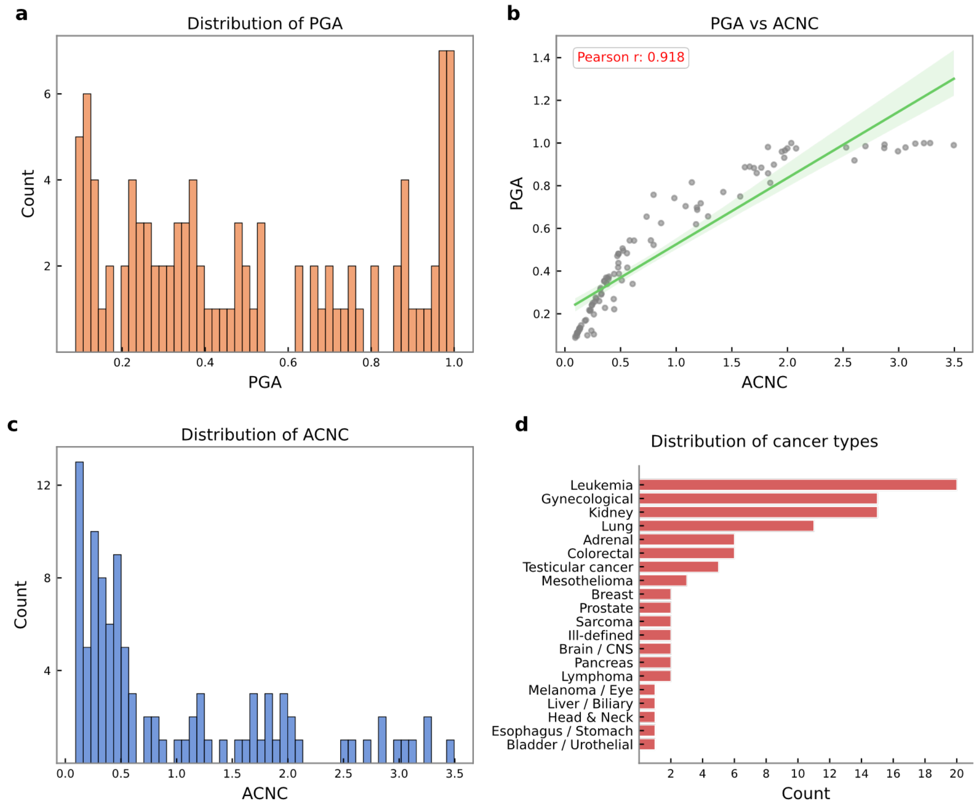


**Fig. S2 Characteristics and diversity of the GDC cancer profiles used for the semi-synthetic cohort. a, c** Histograms showing the distribution of two metrics of genomic complexity across the 100 cancer CNPs randomly selected from the GDC for the in silico spike-in experiments. The cohort includes a broad range of genomic instability, from genomically quiet tumors with low PGA to highly chaotic or whole-genome-doubled genomes with high PGA. **b** Correlation between PGA and ACNC (Pearson's $r = 0.918$) as measurements of genomic complexity. **d** Distribution of cancer types within the GDC cohort. The selection includes diverse cancer types to demonstrate generalizability across real-world CNA patterns.

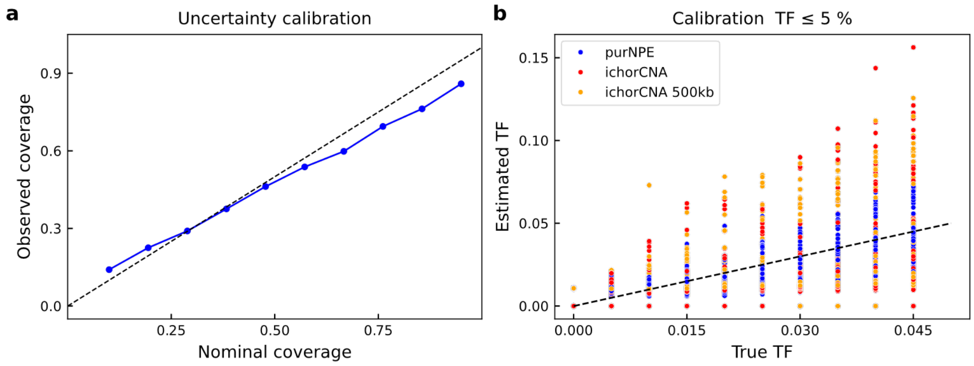


**Fig. S3 Calibration of purNPE Bayesian uncertainty estimates and performance in the low-TF regime.** **a** Uncertainty calibration plot for purNPE evaluated on the semi-synthetic cohort. Observed coverage tracks nominal coverage, shown as the dashed expected-frequency line, demonstrating that posterior distributions are calibrated. **b** Comparison of estimated versus true TF for all methods in the low-TF regime (true TF $\leq 5\%$). purNPE estimates are tightly distributed around the ground-truth $y = x$ line. Estimates from ichorCNA and ichorCNA 500kb exhibit higher variance and systematic underestimation.

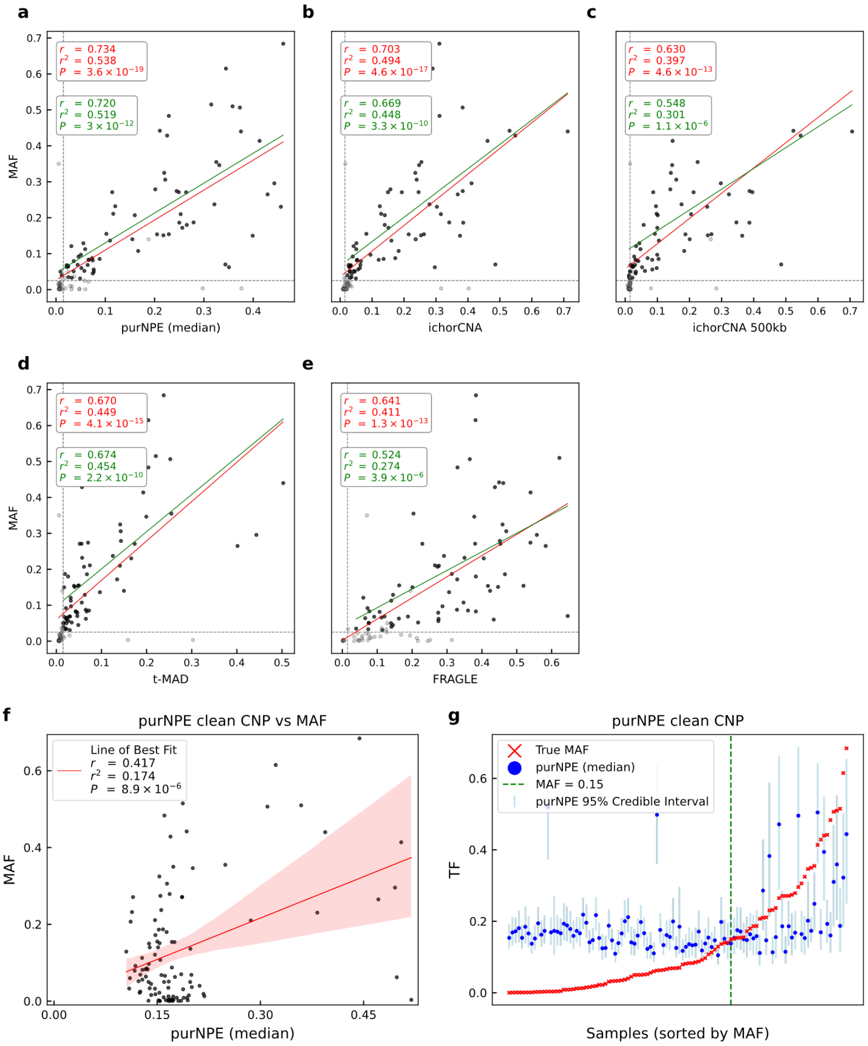


**Fig. S4 Comparison of TF estimators against an orthogonal MAF proxy.** Scatter plots compare TF estimates from **a** purNPE, **b** ichorCNA, **c** ichorCNA 500kb, **d** t-MAD, and **e** FRAGLE against MAF from the Mouliere et al. clinical cohort. To allow direct comparison with the original study, two linear regressions were performed for each method: one across all samples and another on a high-signal subset, defined as samples with MAF $> 0.025$ and t-MAD $> 0.015$. purNPE demonstrates the highest overall correlation (Pearson's $r = 0.734$) across all samples. **f** Demonstration of model failure on the clinical cohort when GNM draws are not considered during training and the model trains on clean CNPs alone; the linear fit is low ($r = 0.417$, $r^2 = 0.174$). **g** On clean profiles, model performance indicates little sensitivity in the low-TF region, while some correlation is observed at high TF ($> 15\%$).

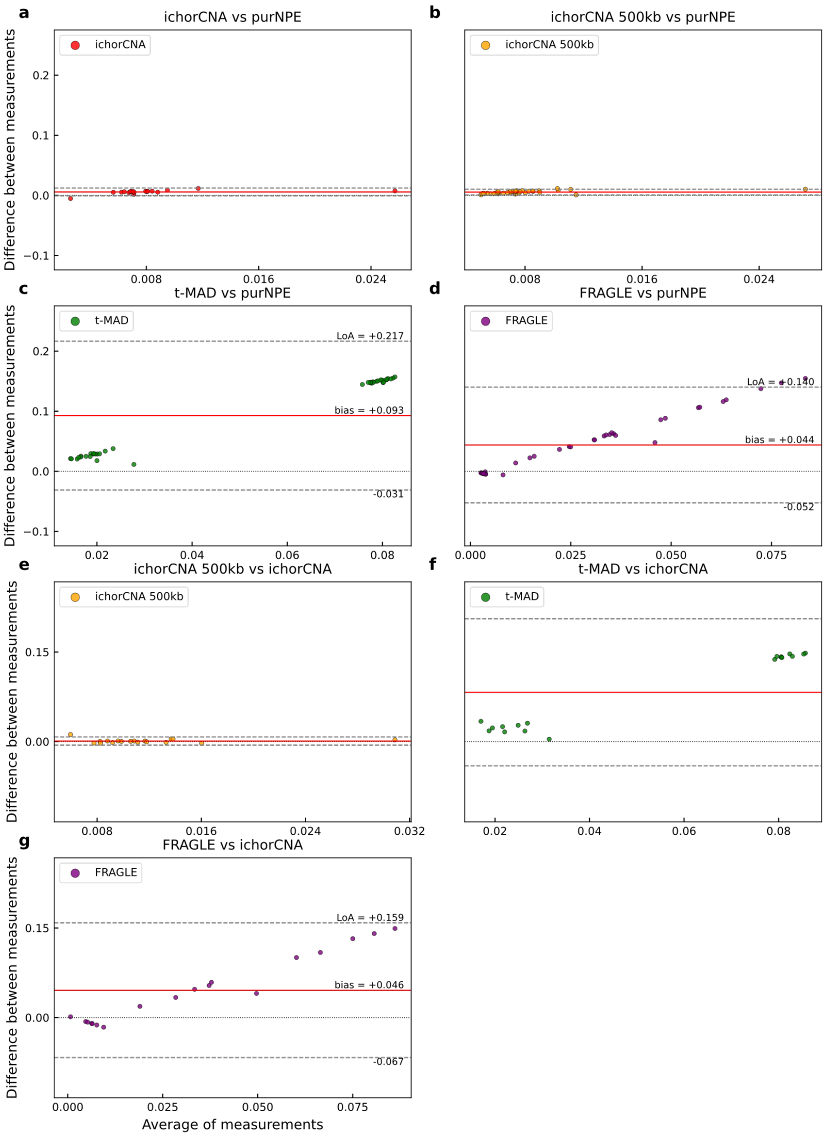


**Fig. S5 Bland–Altman plots for pairwise inter-method agreement on clinical cohorts.** Bland–Altman plots show agreement between purNPE, used as the reference method, and **a** ichorCNA, **b** ichorCNA 500kb, **c** t-MAD, and **d** FRAGLE across two independent clinical cohorts. Each point represents one sample; the $x$-axis shows the average of the two measurements and the $y$-axis shows their difference. The solid red line indicates the mean difference (bias), and dashed grey lines represent the 95% limits of agreement (LoA). **a, b** purNPE shows high concordance with both ichorCNA modes, with very low bias ($\approx +0.5\%$) and tight LoA under the evaluated conditions. **c, d** In contrast, purNPE shows systematic disagreement with methods based on orthogonal signals, with a large positive bias for t-MAD (+9.3%) and FRAGLE (+4.4%). **e–g** Analogous comparisons using ichorCNA.

**Table S1** Detailed architecture of the purNPE inference network.

| Layer / Block | Description | Key parameters | Output shape |
|---|---|---|---|
| Initial convolution | A 1D convolutional layer to create initial feature maps. | `out_channels=128`; kernel = 7 | 4,784 |
| Downsampling block 1 | 1D convolution with stride 2 to halve sequence length. | `out_channels=128`; kernel = 7; stride = 2 | 2,391 |
| Downsampling block 2 | 1D convolution with stride 2. | `out_channels=128`; kernel = 7; stride = 2 | 1,196 |
| Downsampling block 3 | 1D convolution with stride 2. | `out_channels=128`; kernel = 7; stride = 2 | 598 |
| Residual group 1 | Three residual blocks to deepen feature extraction. | channels = 128; `n_blocks=3` | 598 |
| Downsampling block 4 | 1D convolution with stride 2. | `out_channels=128`; kernel = 7; stride = 2 | 299 |
| Residual group 2 | Two residual blocks. | channels = 128; `n_blocks=2` | 299 |
| Residual group 3 | Four residual blocks. | channels = 128; `n_blocks=4` | 299 |
| Global pooling and head | Collapses the sequence and generates the final embedding. | Dropout ($p$ = 0.05); adaptive average pooling; linear layer with `out_features=128` | 1 |
| Output embedding | A 128-dimensional summary vector representing the input CNP. | – | 128-dim. vector |

The purNPE inference network is an NPE composed of a deep one-dimensional ResNet1D encoder and an NSF density estimator. The input is a 500-kb binned, $\log_2$-transformed, corrected read-depth profile of length 4,784 bins. The ResNet1D encoder has approximately 2.75 million trainable parameters. The NSF component consists of five masked autoencoder for distribution estimation blocks, each with two hidden layers of 128 units and ReLU activation.

***Table S2. Source and metadata for the 100 GDC cancer profiles used in the in silico validation cohort.***

This table provides file identifiers and key metadata for the 100 copy-number segment files downloaded from the publicly available GDC data portal. These files provided the ground-truth cancer CNPs used to generate the in silico spike-in benchmark cohort analyzed in Figs. 2, 3, and 5 and Supplementary Figs. S1 and S2. The full, unabridged table is available as a supplementary data file.

***Table S3. Per-sample TF estimates for all methods on the Mouliere et al. clinical cohort.***

This table provides raw per-sample TF estimates from purNPE, ichorCNA, ichorCNA 500kb, t-MAD, and FRAGLE for each of the 106 clinical samples from the Mouliere et al. cohort. The MAF column represents the orthogonal biological proxy used for comparison. This is the source data for analyses presented in Fig. 4, Supplementary Fig. S4, and Table 1. The full, unabridged table is available as a supplementary data file.

***Table S4. Per-sample TF estimates for all methods on independent clinical cohorts.***

This table provides raw per-sample TF estimates from purNPE and all competitor methods for the independent clinical cohort samples used in the study. These values support the inter-method concordance analyses and, for control samples, represent the baseline signal or noise floor used in the semi-synthetic LoD and ROC analyses. The ichorCNA column contains NaN values for this mode because control samples were used to construct the cohort-matched PoN and were therefore part of the reference rather than evaluated samples. The full, unabridged table is available as a supplementary data file.

***Table S5. Metadata for cancer-free control cfDNA samples.***

This table provides metadata for the cancer-free control cfDNA sequencing samples used to train the GNM. The full, unabridged table is available as a supplementary data file.

***Table S6. Metadata for the Mouliere et al. clinical cohort.***

This table provides metadata for the Mouliere et al. pan-cancer cohort sequencing. The full, unabridged table is available as a supplementary data file.

***Table S7. Metadata for the independent PDAC clinical cohort.***

This table provides metadata for the PDAC cfDNA sequencing cohort. The full, unabridged table is available as a supplementary data file.

***Table S8. Metadata for the independent breast cancer clinical cohort.***

This table provides metadata for the breast cancer cfDNA sequencing cohort. The full, unabridged table is available as a supplementary data file.

### Extended methods

#### Sequencing data pre-processing and read-depth profile generation

All aligned sequencing data in BAM format, including clinical samples and cancer-free controls, underwent a standardized pre-processing pipeline to generate corrected read-depth profiles for copy-number analysis. This process began from aligned reads represented in the Sequence Alignment/Map format and related BAM files [1]. Genome-wide read counts were quantified by binning the genome into non-overlapping windows of fixed size. The `readCounter` utility from HMMcopy was used for this task [2]. To ensure that only uniquely and confidently mapped reads were included, low-quality or multiply mapped reads were filtered before read-count aggregation.

To account for known systematic biases in WGS data that affect read depth, we either generated or downloaded corresponding GC-content and mappability profiles for each required bin size. The `gcCounter` utility was used to calculate the GC fraction within each bin, while mappability tracks were used to represent the uniquely mappable fraction of each genomic interval. The raw read counts for each sample were then corrected for GC and mappability biases. The resulting values were $\log_2$ transformed to produce the final corrected read-depth profile used as input for all downstream analyses. The overall sequencing-processing strategy follows standardized principles for scalable and reproducible DNA-sequencing analysis [3].

#### Competitor methods

Analysis of ichorCNA was performed using ichorCNA v0.5.0 [4]. Default parameters were used with all autosomal chromosomes. The following changes to default parameters were used:

```
centromere='GRCh38.GCA_000001405.2_centromere_acen.txt'
scStates='', repTimeWig='', ploidy=2, includeHOMD=FALSE
estimateNormal=TRUE, estimatePloidy=TRUE, altFracThreshold=0.05
genomeBuild='hg38', genomeStyle='UCSC'
fracReadsInChrYForMale=0.001, txnE=0.9999, txnStrength=10000
normal='c(0.5,0.6,0.7,0.8,0.9,0.95,0.99,0.995,0.999)'
```

A panel of normals (PoN; ichorCNA parameter `normal_panel`) was used for the ichorCNA mode with the respective matching samples. Mappability and GC files for 50-kb and 500-kb bins were taken from the ichorCNA resource. For the ichorCNA 500kb mode, the default PoN was used to represent execution without a study-specific PoN. The t-MAD metric served as a simple, non-model-based proxy for the overall magnitude of CNAs. For re-analysis of the Mouliere et al. cohort [5], t-MAD values were used directly as reported in the original publication. For the two independent clinical cohorts, t-MAD was calculated from shallow WGS data using the same definition. FRAGLE v1.1 predicts TF by analyzing the density distribution of cfDNA fragment lengths [6]. Input BAM files were provided to the trained model as described by the authors in the FRAGLE software repository [7]. The PDAC and breast cancer cohorts were processed using the same input-generation strategy.

#### Semi-synthetic benchmark design and generation

To assess model performance against known ground truth, we generated a semi-synthetic in silico spike-in benchmark. This process involved projecting high-resolution real tumor CNPs onto a fixed-bin format and computationally mixing them with an unseen cancer-free cfDNA noise profile. We sourced 100 diverse, allele-specific absolute copy-number segment files from ASCAT profiles available through the GDC data portal. These files represent real tumor genomic architectures, but are provided as segmented profiles rather than raw sequencing reads. For the background noise component, we used the corrected $\log_2$ read-depth profile from a single randomly selected cancer-free control cfDNA sample from the PDAC cohort. This sample was strictly held out from GNM training.

Each of the 100 curated GDC profiles was computationally mixed with the unseen control noise profile across 30 discrete TFs ranging from 0.5% to 15% in 0.5% increments. This procedure produced a benchmark dataset of 3,000 genomes. Each resulting profile has a known ground-truth tumor architecture, a precise ground-truth TF, and an unseen sequencing-noise pattern.

#### In silico LoD and sensitivity assessment

To assess model performance in the low-TF regime, a dedicated in silico cohort was constructed by computationally spiking the GDC profiles into cancer-free controls from the PDAC and breast cancer cohorts. Each of the 100 GDC tumor profiles was spiked into each of the 21 control profiles. This procedure was repeated across a dilution series with expected TFs of 0.5%, 1.0%, 2.0%, 3.0%, 4.0%, and 4.9%. The original 21 cancer-free profiles served as the baseline 0% TF control group.

Analytical sensitivity was estimated according to CLSI EP17-A2 guidelines. This involved estimating the LoB and LoD from a large set of semi-synthetic replicates. A set of 900 blank replicates was generated from non-spiked profiles (0% TF). A corresponding set of 900 low-level replicates was generated by spiking a 1% TF signal. The LoB was defined as the 95th percentile of the blank-sample distribution, and the LoD was calculated as

$$\mathrm{LoD} = \mathrm{LoB} + 1.645 \cdot \sigma_{\mathrm{low}}, \tag{1}$$

where $\sigma_{\mathrm{low}}$ is the standard deviation of the low-level replicates.

#### Patient recruitment and sample collection

This study was conducted in accordance with the ethical standards of the institutional research committees. The study was approved by the Helsinki Committee of the Sheba Medical Center (approval number 9534-22-SMC) and the Helsinki Committee of the Tel Aviv Sourasky Medical Center (approval number 0569-16-TLV). Informed consent was obtained from all participants prior to sample collection.

The PDAC cohort included patients at stages I to IV, with samples collected at various clinical time points (e.g., pre-adjuvant, on-treatment, post-resection). The breast cancer cohort consisted of female patients diagnosed predominantly with infiltrating ductal carcinoma (Stages I–II, based on pTNM staging) with varying hormone receptor statuses (ER, PR, Her-2). The control group for both cohorts included individuals with no known malignancy or, for the breast cancer cohort, individuals with benign conditions such as fibroadenoma.

Across both cohorts, participants ranged in age from 21 to 80 years. Blood was collected using ethylenediaminetetraacetic acid (EDTA) tubes for all participants, except for one healthy control sample collected in a Streck tube.

#### Plasma separation and cfDNA extraction

For the PDAC cohort, whole blood samples were processed promptly in-house to isolate plasma. The procedure involved an initial centrifugation at 1,500 $\times g$ for 10 minutes at 4°C using Lymphoprep density-gradient medium, followed by an additional centrifugation step to remove cellular debris. The resulting plasma was aliquoted and stored at −80°C until cfDNA extraction. Plasma from patients with breast cancer and benign tumors was obtained from the TLV BioBank, Tel Aviv Sourasky Medical Center, Tel Aviv, Israel. For this cohort, whole blood samples were collected in EDTA tubes, centrifuged, and stored according to the BioBank protocol.

Subsequently, cfDNA was extracted using the QIAamp Circulating Nucleic Acid Kit (Qiagen), following the manufacturer's protocol. The starting plasma volume varied between cohorts, with 2–4 mL used for PDAC samples and a smaller available volume used for the breast cancer cohort according to sample availability.

##### cfDNA quality assessment

The quality and fragment-size distribution of extracted cfDNA were evaluated using the Agilent 4200 TapeStation System with the Cell-free DNA ScreenTape assay. This platform allows determination of the proportion of cfDNA fragments within the expected mono-nucleosomal range and detection of potential high-molecular-weight contamination. Samples passing quality assessment were used for library preparation and sequencing.

##### Library preparation and whole-genome sequencing

Sequencing libraries were prepared and sequenced for the two clinical cohorts at different facilities using distinct protocols. All resulting raw sequencing data in FASTQ format were then processed through the same standardized downstream analysis workflow. For the breast cancer cohort ($n = 19$), following DNA extraction, libraries were generated using a standard ligation-based workflow for whole-genome resequencing. Prepared libraries were sequenced on an Illumina platform to generate $2 \times 143$ bp paired-end reads. The sequencing run yielded a mean of 22.1 Gb of data per sample.

For the PDAC cohort ($n = 21$), sequencing libraries were prepared from cfDNA using the Accel-NGS 2S PCR-Free kit, which is optimized for low-input samples and minimizes amplification bias. Libraries were subsequently sequenced on an Illumina NovaSeq X Plus platform to generate $2 \times 151$ bp paired-end reads. This process targeted an output of 10 Gb of data per sample.

[3] Hanssen F, Garcia MU, Folkersen L, Pedersen AS, Lescai F, Jodoin S, et al. Scalable and efficient DNA sequencing analysis on different compute infrastructures

aiding variant discovery. NAR Genomics and Bioinformatics. 2024;6(2):lqae031. https://doi.org/10.1093/nargab/lqae031.

[4] Adalsteinsson VA, Ha G, Freeman SS, Choudhury AD, Stover DG, Parsons HA, et al. Scalable whole-exome sequencing of cell-free DNA reveals high concordance with metastatic tumors. Nature Communications. 2017;8(1):1324. https://doi.org/10.1038/s41467-017-00965-y.

[5] Mouliere F, Chandrananda D, Piskorz AM, Moore EK, Morris J, Ahlborn LB, et al. Enhanced detection of circulating tumor DNA by fragment size analysis. Science Translational Medicine. 2018;10(466):eaat4921. https://doi.org/10.1126/scitranslmed.aat4921.

[6] Zhu G, Rahman CR, Getty V, Odinokov D, Baruah P, Carrié H, et al. A deep-learning model for quantifying circulating tumour DNA from the density distribution of DNA-fragment lengths. Nature Biomedical Engineering. 2025;9(3):307–319. https://doi.org/10.1038/s41551-025-01370-3.

[7] Skandalis Lab. FRAGLE software repository [web page]. GitHub; 2026 [cited 18 May 2026]. Available from: https://github.com/skandlab/FRAGLE.
